## Supplementary figures and text for "Neuronal Variability Reflects Probabilistic Inference Tuned to Natural Image Statistics"

### Supplementary Information – Neuronal Variability Reflects Probabilistic Inference Tuned to Natural Image Statistic

#### Supplementary Figures

S1 Additive Gaussian mixer model . . . . . 2
S2 Training the GSM with different statistics . . . . . 4
S3 Data, mean-matching controls . . . . . 5
S4 Data, relation between surround modulation of response and of FF . . . . . 6
S5 GSM model, estimate of the mixer variable for size tuning and orientation tuning . . . . . 7
S6 Data, size tuning for grating stimuli . . . . . 8
S7 Data, size tuning experiments divided by RF size . . . . . 9
S8 Reduced GSM model, centered versus offset stimulus . . . . . 10
S9 Relation between surround modulation of rate and of FF . . . . . 11
S10 Response of GSM model for different levels of additive noise . . . . . 12
S11 Contrast tuning and orientation tuning in the GSM model . . . . . 13
S12 Size tuning and surround orientation tuning, GSM with rectified-supralinear activation . . . . . 14

### Supplementary Text


1 Statistics of the GSM mixer in the low-noise approximation 15

2 Statistics of one latent variable in the low noise approximation 16

3 Statistics of the latent feature for the special case of Rayleigh mixer prior 17

4 Approximate posteriors for mixer and latent variable 18

5 Conversion from latent variables to spike counts 19

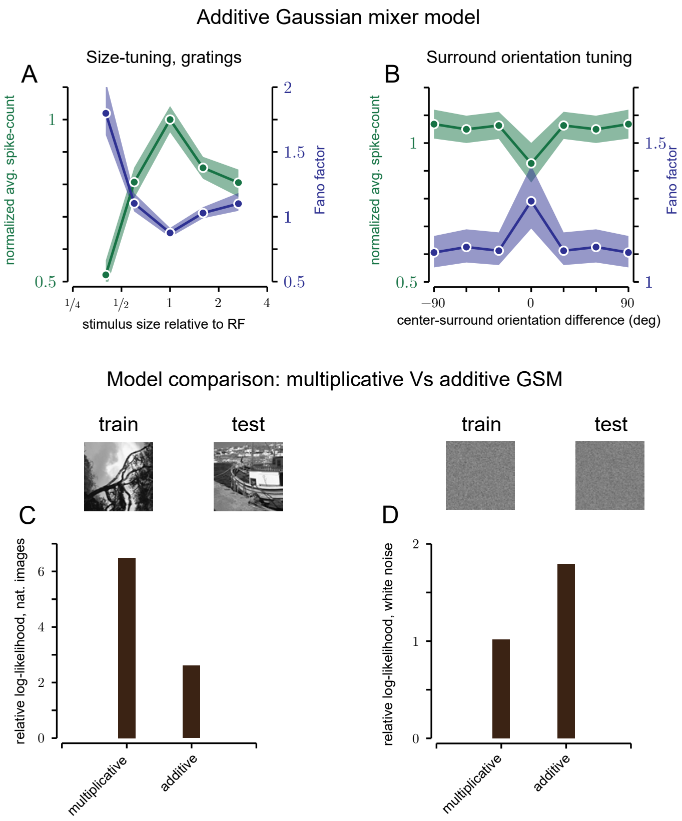


**Figure S1:** **A,B**. Alternative model with additive rather than multiplicative mixer term. When considering mean spike counts, the model produces qualitatively similar behavior to the multiplicative model for size-tuning (panel **A**, green) and surround orientation tuning (panel **B**, green). However, in the additive formulation the variance is constant and does not depend on the input, therefore the Fano factor is simply a rescaled version of the inverse of the mean (blue lines in both panels), in contrast with the multiplicative GSM and with experimental findings (Fig. 2D-F and Fig. 3A-C main text). **C,D**. Model comparison between additive and multiplicative GSM for different image statistics. The plot reports the average log-likelihood over a set of 10,000 test image patches for a multiplicative and an additive GSM, both without additive noise. The measure is normalized so that 0 corresponds to the log-likelihood for a null model where each element of **x** is an independent Normal. As expected, the multiplicative version of the GSM is better adapted to natural image statistics Wainwright. White noise statistics, instead, can be captured by a multivariate Normal distributions, therefore the additive model, still Normal in nature, performs better than the multiplicative.

**Methods.** For these figures only, the generative model takes the form $\mathbf{x} = \nu \,.\!+ \mathbf{g} + \boldsymbol{\eta}$ . The input filters are the same used for the GSM, however here the mixer $\nu$ is a scalar *additive* term, with $.+$ indicating the scalar-vector sum. Since a positive-only mixer might bias the posterior $P(\mathbf{x})$, we chose the prior $\nu \sim \mathcal{N}(0, \sigma_\nu)$ for the mixer. To train the model, the mixer was optimized to match the statistics of mean $(\mathbf{x})$, over natural image patches. The covariance structure of the features was trained using expectation maximization over 10,000 natural image patches, assuming a noiseless model. We then added a noise term with the same structure and scaling used for the GSM. Finally, to convert the latent feature variable into a spike count, we used: $r = \alpha\,(g_{1+} + \beta)$, with $\beta$ sufficiently high to avoid negative terms. The phase of the input gratings was also regulated so that the filters output would be positive. For panels **C,D** we considered models without additive noise. We drew the training patches from 80% of the image dataset (which comprises 500 full scale images in total), and the test patches from the remaining 20%. Lastly, the models used in **D** were optimized using white noise patches instead of natural patches.

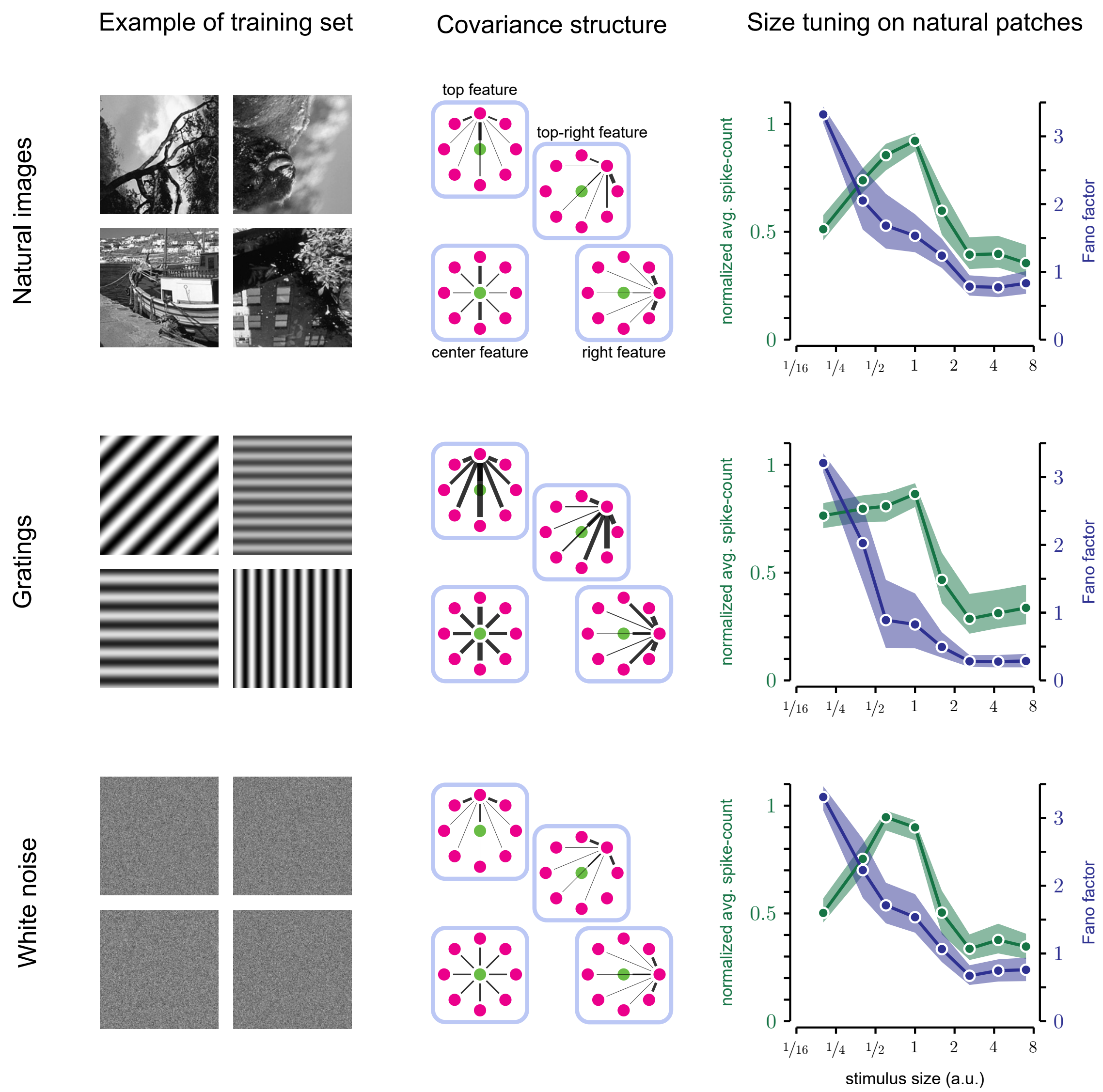


*figure caption on next page*

**Figure S2:** GSM model trained on different input statistics. **Left column.** Examples of training images. We used 10,000 random patches of natural scenes (top row), uniform gratings with random orientation, frequency and contrast (middle row), or white noise (bottom row). To compare the effects of different training sets quantitatively, we considered the noise level as the relative scaling between the average of the diagonal terms of $C_g$ and of $C_{\text{noise}}$, and not, as before, as absolute scaling of $C_{\text{noise}}$. For this reason the model outputs (right column) are slightly different, even when training on natural images.

**Center column.** covariance structure after training. Circles represent the localized GSM features with vertical orientation, and edges indicate the positive correlations between them, corresponding to the off-diagonal elements of $C_g$. Features that are very positively correlated share a thicker edge, while independent features are not connected. Within each panel, each filter bank, enclosed by a square, represents the correlation between one filter and all others. Only three of the eight surround filters are shown, as the structure is symmetric for 90° rotations, reflecting that input images were randomly rotated by $90°$,$-90°$ and $180°$.

**Right column.** Tuning of spike-count mean and FF for natural image patches varying in size. Inputs are the same used in Fig. 2D, main text. The three models produce qualitatively similar curves. Note however that in an GSM-based model of neural populations, training on natural images versus white noise would lead to qualitatively different predictions. For instance, pairwise noise correlations would depend strongly on the geometry of the receptive fields for the GSM trained on natural images, but depend only on the distance between receptive fields for the GSM trained on white noise. This, and related predictions, could be tested in the future with new, targeted experiments.

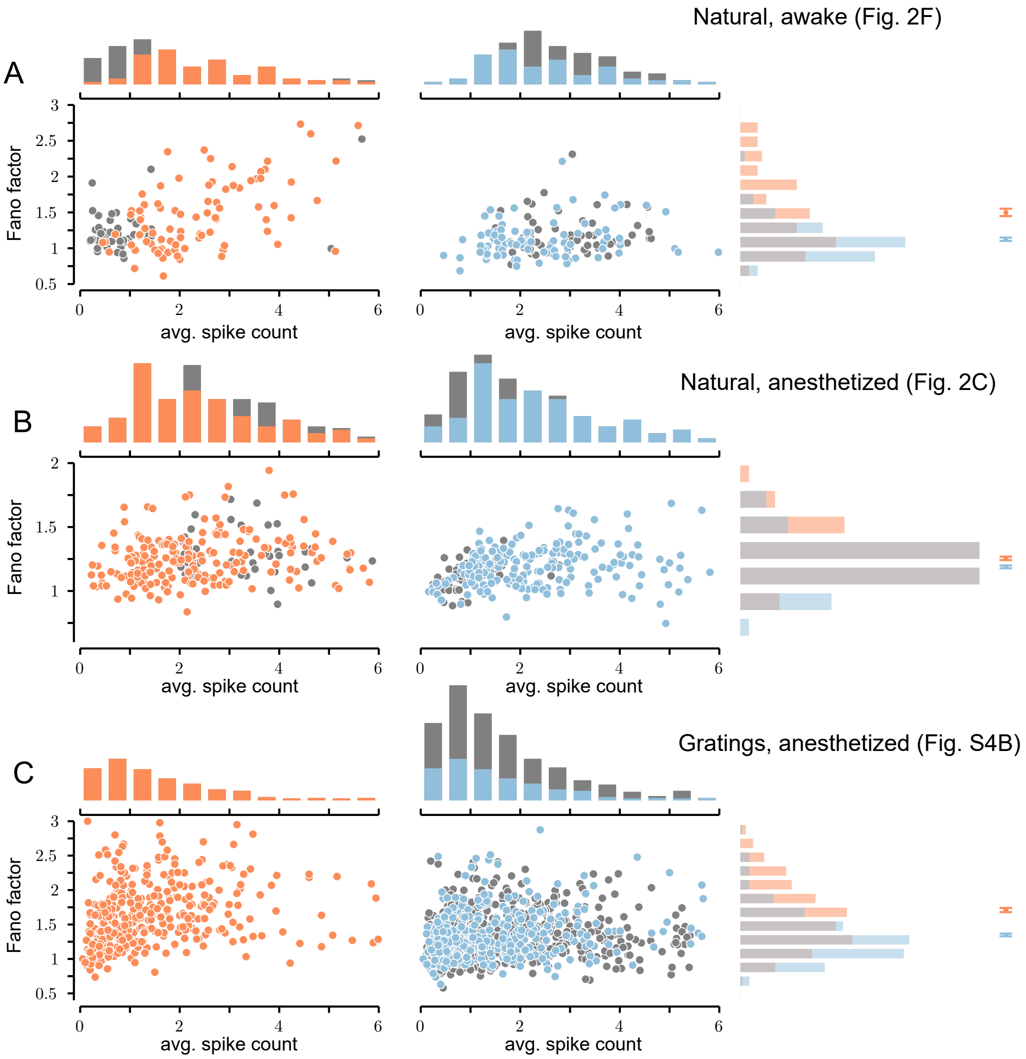


**Figure S3:** Mean-matching for the size tuning experiments. Procedure: we first split the data based on whether stimulus size was smaller or larger than the RFs (left and right panels of each figure, respectively). We then computed the spike-count mean and FF for each combination of size/neuron (individual dots in the scatter plots). For each condition, we then subsampled the size/neuron elements so that the spike-count histograms were identical in the two conditions (orange and blue histograms above). Lastly, for those selected points, we considered the Fano factor distributions (histograms on the right) and the population mean (bars are 95% c.i., computed by bootstrapping). In all cases the Fano factors were significantly lower for larger stimuli.

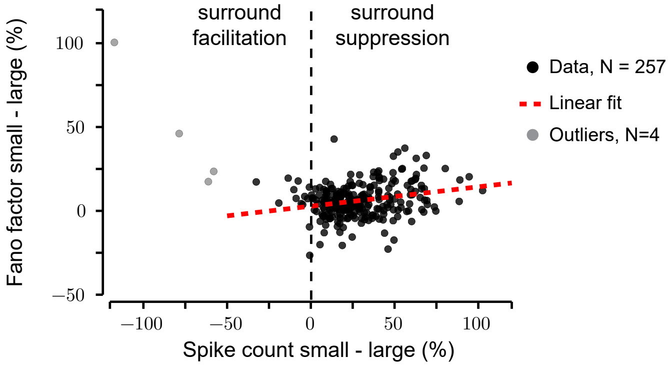


**Figure S4:** This plot refers to the same experiment shown in Fig. 2C of main text: stimuli consisted of small and large patches of natural images, and neuronal responses were measured in V1 of anesthetized macaques. The plot compares the surround suppression score of the spike-count mean and FF. Positive scores denote a reduction from small to large images, according to: $100 \cdot \frac{Z_{\text{small}} - Z_{\text{large}}}{0.5(Z_{\text{small}} + Z_{\text{large}})}$, where $Z$ refers to either the spike-count mean or to the FF for each condition (see Methods).Each point represents one neuron; 4 neurons (gray symbols) had a spike-count score below $-50\%$, i.e. unusually strong surround facilitation that suggests poor centering; these outliers have been excluded from further analysis (but are included in the main Fig. 2C and related text). The remaining neurons showed a significant correlation between surround suppression of spike counts and of FF (Pearson corr. $0.25$, $p < 10^{-4}$).

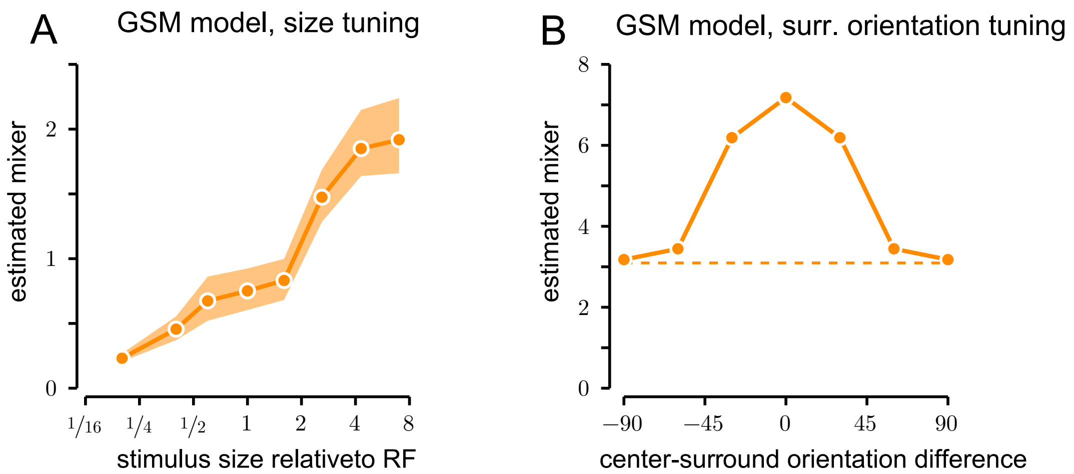


**Figure S5:** GSM model, numerical estimate of the mixer posterior. **A.** (Corresponding to Fig. 2D of the main text) Estimate of the global mixer for natural image patches of increasing size. The mixer grows monotonically, which causes the monotonic reduction of FF appearing in Fig. 2D. **B.** (Corresponding to Fig. 3A of the main text) Estimate of the global mixer for a surround of varying orientation (relative to the center stimulus). Dashed line: center stimulus only. The mixer increases when the surround is matched (parallel) to the center, corresponding to the drop in FF observed in Fig. 3A.

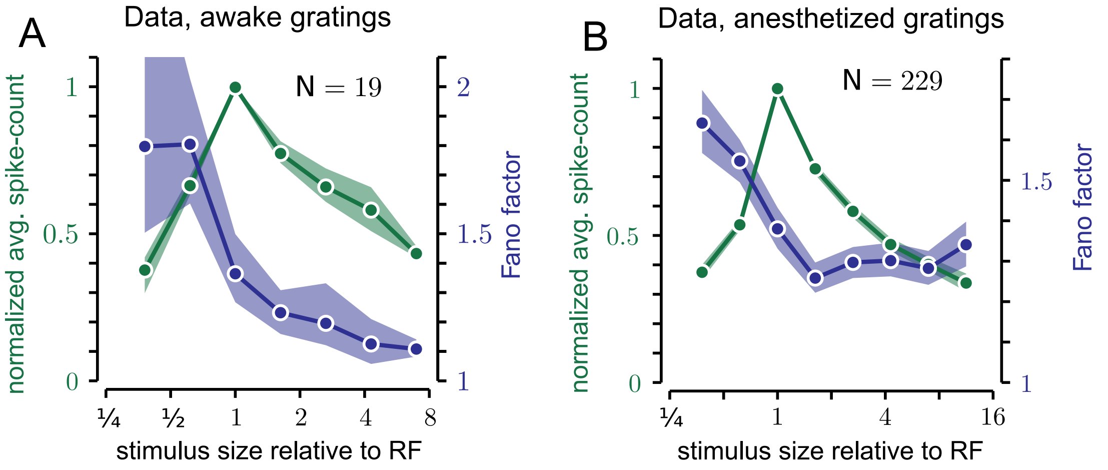


**Figure S6:** Data, population size turning for grating stimuli. **A**. One awake fixating monkey; **B**. Three anesthetized monkeys; same conventions as in Figure 2F main text.

**Methods:** spike counts at each trial were computed as specified in Methods, main text. Spike-count means and FFs were first calculated across trials separately for each stimulus condition (spatial phase in anesthetized experiment, and size and orientation in both). Then all conditions except size were averaged for each neuron, and means were normalized by RF size. Lastly, the population average was computed across all neurons. Error bars represent the 68% c.i.

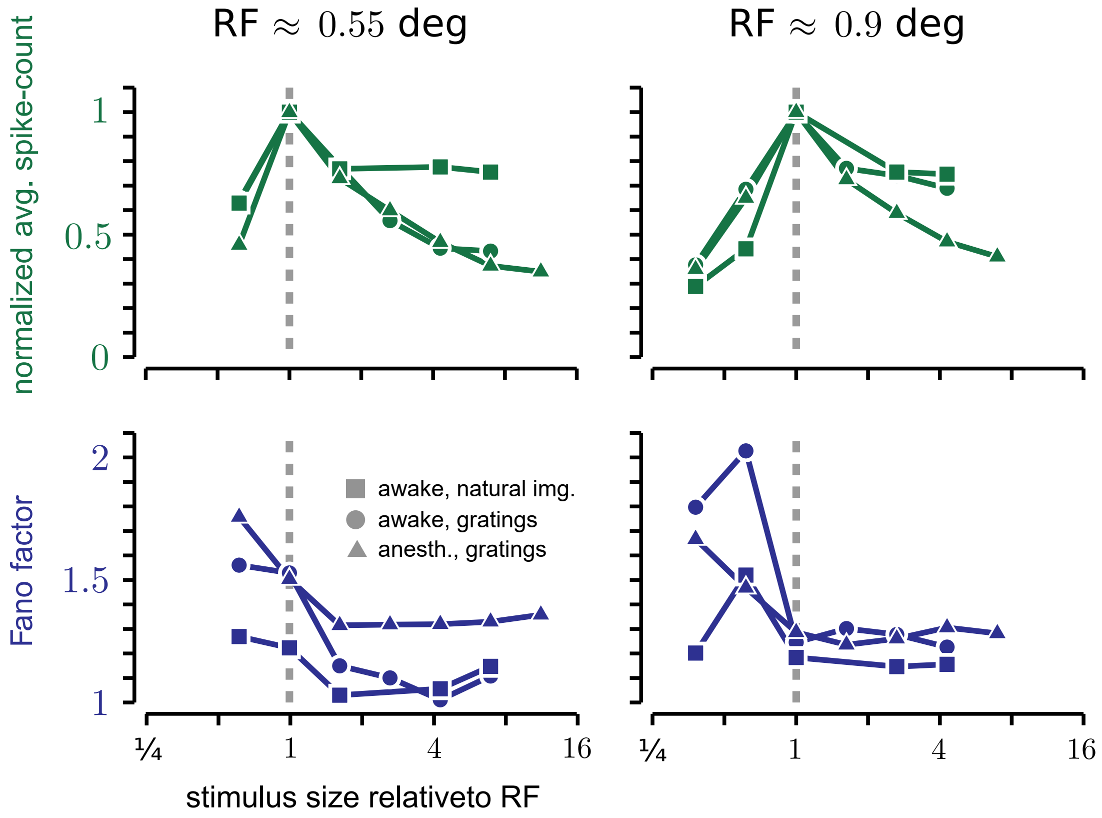


**Figure S7:** Population averages for size tuning experiments (experiments represented by distinct symbols), where populations have been split based on whether the RF size was $\approx 0.55$ (left column) or $\approx 0.9$ (right columns). Surround suppression of FF was approximately monotonic for neurons with smaller RF (left panels), whereas neurons with larger RFs (right panels) tend to have a decrease in FF for very small stimuli, and show a weaker suppression of FF by surround.

**Notes:** the RF size was computed separately across stimulus parameters such as image identity (for natural images), grating orientation, and spatial phase. Some neurons therefore appear in both categories, but with different stimuli. Populations have been split as follows: natural image patches, awake, 86 neurons in total , $26\%$ had RF $= 0.55$ deg ; gratings, awake, 19 neurons in total, $42\%$ with RF $= 0.55$ deg; gratings, anesthetized 229 neurons in total, $46\%$ with RF $= 0.55$ deg.

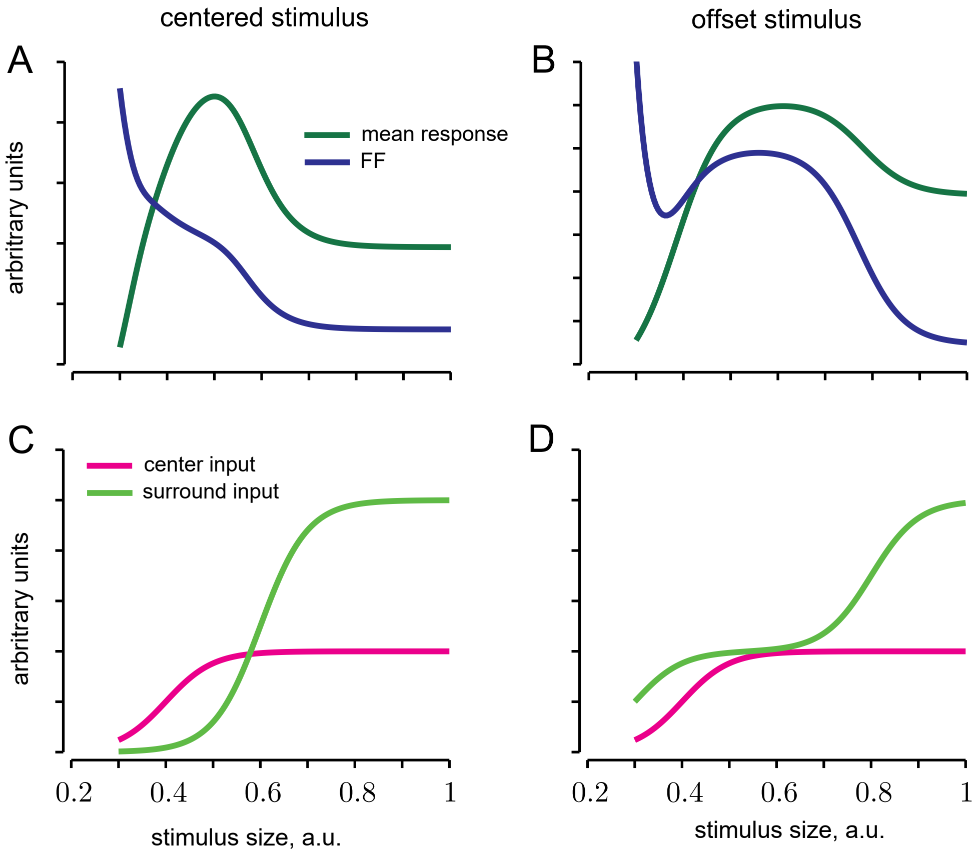


**Figure S8:** Responses in a 2D GSM model for a well-centered (left) and an offset stimulus (right). **A**. When the stimulus was well centered, the mean (dark green) associated to the central feature had surround suppression, and the FF (blue) decreased monotonically. **B**. When the stimulus was offset, instead, the mean response of the model peaked at a larger size (dark green), and the FF was non-monotonic (blue).

**C-D**. Inputs used in the two conditions: in the first case (panel C) center and surround were activated in sequence (magenta and light green line, respectively). Instead the offset stimulus (panel D) first activated part of the surround, but not the center, causing the initial drop in FF, then activated the center, and finally the remaining surround areas.

**Methods:** We opted for a 2D GSM model to remove possible confounders, such as dependencies due to the filter covariance structure. The two dimensions represent, respectively, the encoded latent variable (the center), and the contextual information (the surround). The model parameters $C_g$ and $C_{\text{noise}}$ (Eq. S1) were both diagonal, with elements $[5, 5]$ and $[0.01, 0.01]$ respectively. The inputs varied parametrically according to size, taking a sigmoidal shape. The center input saturated at $1$, whereas the surround input reached a level of $2.5$. For simplicity, we considered the input as only positive, and directly identified the spike counts with the latent posterior distribution. Mean and FF of the model response were calculated semi-analytically by numerical integration.

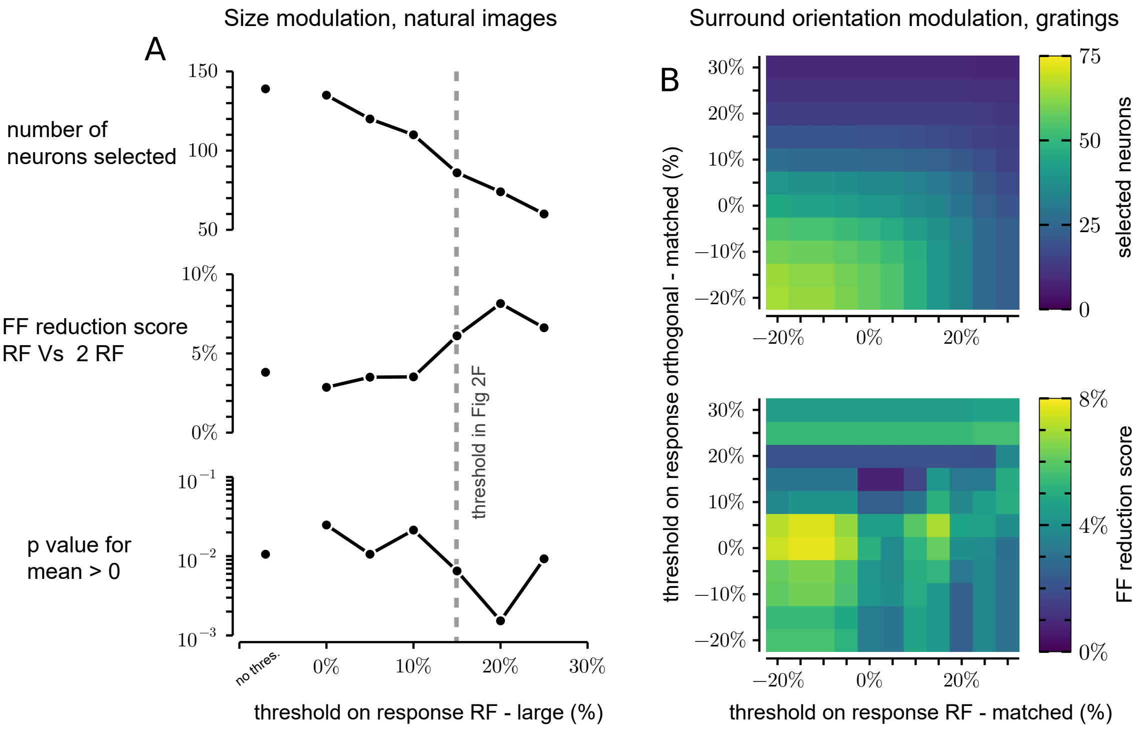


**Figure S9: A**. Robustness to neuron selection criteria for the size tuning experiment with natural images patches (main text, Fig. 2F). Neurons have been selected based on a spike-count score given by: $100 \cdot \frac{r_\alpha - r_\beta}{(r_\alpha + r_\beta)^{1/2}}$, where $r$ indicates the spike-count mean across repetitions, and $\alpha$ and $\beta$ refer to RF size versus large stimuli. The population average of the FF was evaluated using the score: $100 \cdot \frac{\mathrm{FF}_\alpha - \mathrm{FF}_\beta}{(\mathrm{FF}_\alpha + \mathrm{FF}_\beta)^{1/2}}$ (main text Methods, Eq. 5), where $\alpha$ and $\beta$ again represent the two sizes. The x axis reports the threshold for the inclusion criterion, the three panels show (from top to bottom), number of selected neurons, population FF reduction score, and the p-value for the distribution of the FF score having positive mean. For Figure 2F we chose a score of $15\%$ (vertical dashed line). **B**. Relation between surround modulation of rate and surround modulation of FF for size tuning experiments (main, Fig. 3B,C). Scores have been computed as in the equations above, comparing center-only with matching surround (size suppression score) or orthogonal surround with matching surround (surround-orientation suppression score). The color maps show the number of selected neurons when both thresholds were imposed (top panel) and the corresponding FF surround orientation suppression score (lower panel). Note that in Fig. 3 of main text we did not impose any threshold, selecting all 71 neurons.

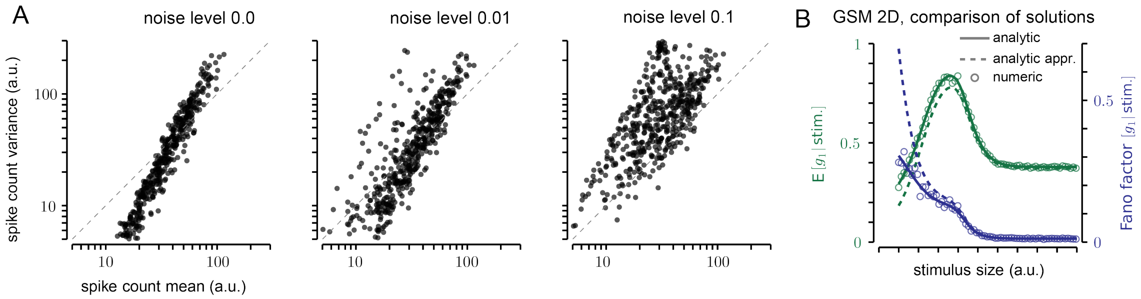


**Figure S10:** **A**. Comparisons of mean-variance plots for GSMs that differ in noise level. The values of $C_{\text{noise}}$ are scaled so that trace$(C_{\text{noise}})/$trace$(C_g) = \gamma$ (see Eq. S1), with noise levels $\gamma = (0,\ 0.01,\ 0.1)$, $0.1$ being the level used in the main text results. Conversion parameter in Eq. S26: $c = 20$. Inputs are natural image patches. Intuitively, when noise is present, it becomes unclear whether a small stimulus is due to a small feature level, or to noise. This results in higher uncertainty for signals that elicit weak responses, therefore the variance increases. **B**. Mean and Fano factor (FF) of the first latent variable in a 2D GSM model with no input noise, as a function of increasing stimulus size. Comparison between full analytical solution (continuous line, see Eq. S15), approximate solution (dashed line, see main text, Eq. 3), and numerical solution (empty circles, see main text, Methods).

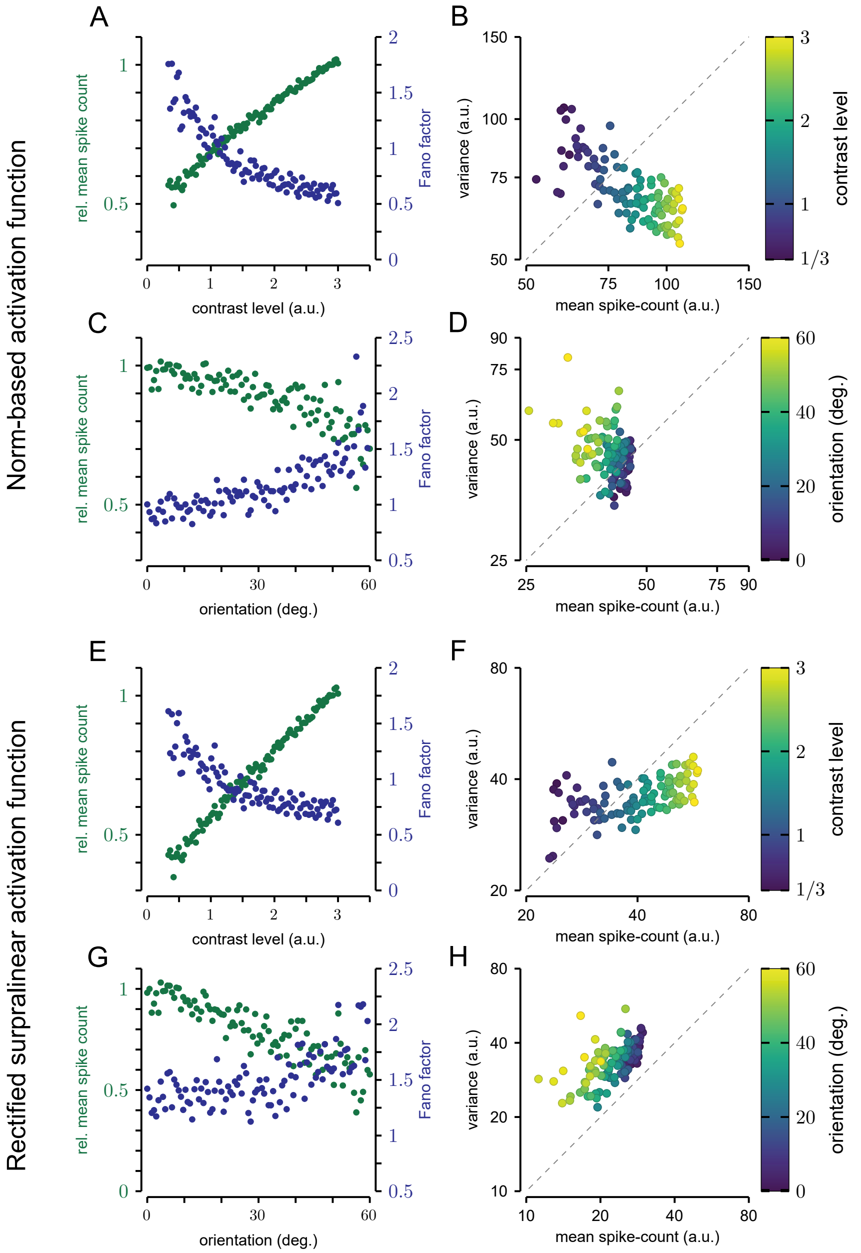


**Figure S11:** **A,B**. and **E,F**. GSM model response for grating stimuli of optimal orientation and varying contrast. **C,D**. and **G,H**. GSM model response for grating stimuli of fixed contrast and varying orientation.

**Methods.** The norm-based activation function converts the latent features $g_{1+}$ and $g_{1-}$ in spike counts according to Eq. S26, with $c = 15$ for contrast tuning and $c = 10$ for orientation tuning; the rectified expansive nonlinearity (Orbán et al. 2016) takes the form $r = \alpha \lfloor g_{1+} + \beta \rfloor_+^\gamma$, where $\lfloor \cdot \rfloor_+$ indicates a rectified linear function, and $\alpha = 3$ , $\beta = 0$ , $\gamma = 1.5$ .

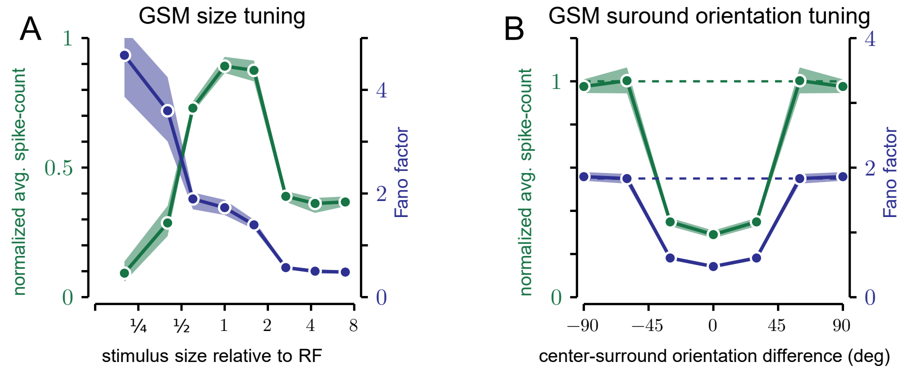


**Figure S12:** GSM model, size tuning for grating stimuli optimized for best response (**A**) and surround orientation tuning (**B**). The function that converts hidden features to spike counts is a rectified expansive nonlinearity (Orbán et al. 2016). The results are qualitatively equivalent to Fig. 2D and Fig. 3A in the main text. The conversion to spike counts follows the form and the parameters described in the caption of Fig. S11.

### Analytical results for the GSM model

In our work we explain the mean and variance of neuronal responses to images in terms of the mean and variance of local latent visual features, computed by probabilistic inference on the GSM model. In Methods Eq. 3,4 we reported analytical approximations for those quantities. Here, we provide the detailed derivations. First, we derive general expressions for the statistics of the global and local latents (Sections 1 and 2, respectively); then we calculate the full distribution of the posterior of the latent variable analytically, and from there, mean, variance and FF (Section 3); then we express these results in an approximate form, easier to interpret (Section 4). Lastly, we show how the latent variable can be transformed into a positive spike count and preserve its properties (Section 5).

For reference, we rewrite here the GSM generative model, as it appears in Eq. 2, main text:

$$\begin{gathered}\mathbf{x} = \nu\,\mathbf{g} + \boldsymbol{\eta} \\ P_g(\mathbf{g}) = \mathcal{N}\left(\mathbf{g}\,;\,0,\,C_g\right) \quad P_\nu(\nu) = \text{Rayleigh}(\nu\,;1) \quad P_\eta(\boldsymbol{\eta}) = \mathcal{N}\left(\boldsymbol{\eta}\,;\,0,\,C_{\text{noise}}\right)\end{gathered} \tag{S1}$$

We refer to $\nu$ as the mixer, to $\mathbf{x}$ as the input, and to $g_1$, i.e. the first element of $\mathbf{g}$, as the local latent variable of interest. Additionally, we use $P_\nu(y)$ to indicate the p.d.f. of the prior on $\nu$ calculated at a generic point $y$. Likewise, $P_g(\mathbf{h})$ indicates the p.d.f. of the Gaussian prior computed at $\mathbf{h}$.

#### 1 Statistics of the GSM mixer in the low-noise approximation

As explained in the main text, the contextual modulatory effects expressed by the GSM are mediated by the global mixer. It is therefore important to understand how the posterior of the mixer behaves as a function of the stimulus, i.e. compute $P(\nu|\mathbf{x})$. Here we consider the low-noise approximation, that is, the case $\|C_{\text{noise}}\| \ll \|C_g\|$. Using Bayes' rule:

$$P(\nu\,|\,\mathbf{x}) = \frac{P(\mathbf{x}\,|\,\nu)\;P_\nu(\nu)}{P(\mathbf{x})} \tag{S2}$$

The first term is the conditional distribution of $\mathbf{x}$ when $\nu$ is known; it can be derived from Eq. S1:

$$P(\mathbf{x}|\nu) = \mathcal{N}\left(\mathbf{x}\,;\,\nu^2\,C_g + C_{\text{noise}}\right) \approx \mathcal{N}\left(\mathbf{x}\,;\,\nu^2\,C_g\right) = \frac{1}{\nu^n}\,\mathcal{N}\left(\frac{\mathbf{x}}{\nu}\,;\,0,\,C_g\right) \tag{S3}$$

where we used the low-noise approximation for the first step, and then we reparametrized the normal distribution, $n$ representing the number of dimensions of $\mathbf{x}$. The denominator of Eq. S2 can then be derived by integrating Eq. S3:

$$P(\mathbf{x}) = \int_0^\infty \mathrm{d}\nu \, P_\nu(\nu) \; P(\mathbf{x}|\,\nu) = \int_0^\infty \mathrm{d}\nu \, P_\nu(\nu) \, \frac{1}{\nu^n} \, \mathcal{N}\left(\frac{\mathbf{x}}{\nu};\, \mathbf{0},\, C_g\right) \tag{S4}$$

If $P_\nu(\nu)$ is Rayleigh, this integral can be solved analytically (Eq. S19). For the time being, we simply define it with the following general notation:

$$\Psi_k\,(\mathbf{x}) := \int_0^\infty \mathrm{d}\nu \, P_\nu(\nu) \; \nu^{(k-n)} \; \mathcal{N}\left(\frac{\mathbf{x}}{\nu}\,;\, 0,\, C_g\right) \tag{S5}$$

Where $n$ is the number of dimensions of $C_g$. Therefore, by definition, $P(\mathbf{x}) = \Psi_0\,(\mathbf{x})$ and:

$$P(\nu|\mathbf{x}) = \frac{1}{\nu^n} \, \frac{\mathcal{N}\left(\frac{\mathbf{x}}{\nu}\,;\, 0,\, C_g\right) \; P_\nu(\nu)}{\Psi_0\,(\mathbf{x})} \tag{S6}$$

Calculating the expectation is straightforward:

$$\mathrm{E}[\nu|\,\mathbf{x}] = \int_0^\infty \mathrm{d}\nu\,\nu \; P(\nu\,|\,\mathbf{x}) = \frac{1}{\Psi_0\,(\mathbf{x})} \int_0^\infty \mathrm{d}\nu \, P_\nu(\nu) \, \frac{1}{\nu^{n-1}} \, \mathcal{N}\left(\frac{\mathbf{x}}{\nu}\,;\, \mathbf{0},\, C_g\right) = \frac{\Psi_1\,(\mathbf{x})}{\Psi_0\,(\mathbf{x})} \tag{S7}$$

An equivalent calculation leads to the second moment, and to the variance:

$$\mathrm{E}[\,\nu^2|\,\mathbf{x}] = \frac{\Psi_2\,(\mathbf{x})}{\Psi_0\,(\mathbf{x})} \quad \text{and} \quad \mathrm{Var}\,[\nu|\,\mathbf{x}] = \frac{\Psi_2(\mathbf{x})}{\Psi_0(\mathbf{x})} - \left(\frac{\Psi_1(\mathbf{x})}{\Psi_0(\mathbf{x})}\right)^2 \tag{S8}$$

In Section 4 we show how, with a specific choice of the prior on $\nu$, we can express these results in an approximate, simpler form, which leads to Eq. 2 in the main text.

#### 2 Statistics of one latent variable in the low noise approximation

Here we are interested in the statistics of selected elements of **g**, given the visual input **x**. Specifically the mean and variance, which we relate to the mean and variance of neuronal activity in the main text. Without loss of generality we denote by $g_1$ the feature of interest and $g_2,\, g_3,\, \ldots\, g_n$ the others. Our goal is to calculate:

$$P(g_1|\,\mathbf{x}) = \frac{1}{P(\mathbf{x})} \int \mathrm{d}g_2 \,\ldots \mathrm{d}g_n \; \mathrm{d}\nu \, \mathrm{d}\boldsymbol{\eta} \; P_g(\mathbf{g}) \; P(\mathbf{x}|\nu,\, \mathbf{g},\, \boldsymbol{\eta}) \; P_\nu(\nu) \; P(\boldsymbol{\eta}) \tag{S9}$$

To find analytic solutions for Eq. S9 we consider once again the low-noise approximation. The $P(\boldsymbol{\eta})$ becomes a Dirac Delta function, centered on zero, and can be integrated out directly. The marginal on $\mathbf{x}$ can also be expressed by a Delta: $P(\mathbf{x}|\,\mathbf{g},\,\nu) \;=\; \delta(\mathbf{x}-\nu\mathbf{g})$. Therefore Eq. S9 becomes:

$$P(g_1|\,\mathbf{x}) = \frac{1}{P(\mathbf{x})}\int \mathrm{d}g_2\,...\,\mathrm{d}g_n\;\mathrm{d}\nu\;P_g(\mathbf{g})\;P_\nu(\nu)\;\delta(x_1-\nu g_1)\,...\,\delta(x_n-\nu g_n) \tag{S10}$$

We can now integrate over the $g_2$, ...$g_n$ latent variables. The argument of each Dirac Delta has a single root, at $g_2 = x_2/v$ ...$g_n = x_n/v$. We can therefore apply the following property: $\int \mathrm{d}x\, f(x)\,\delta(g(x)) = f(x_0)/|g'(x_0)|$, and obtain:

$$P(g_1|\,\mathbf{x}) = \frac{1}{P(\mathbf{x})}\int \mathrm{d}\nu\;\frac{1}{\nu^{n-1}}\;P_g\left(g_1,\;\frac{x_2}{\nu},\;...\;\frac{x_n}{\nu}\right)\;P_\nu(\nu)\;\delta(x_1-\nu g_1) \tag{S11}$$

For the full $g_1$ distribution, we integrate over $\nu$. This is equivalent to the substitution $\nu = x_1/g_1$, and to divide by the norm of the argument of the Delta. Finally we replace $P(\mathbf{x})$ following the definition of Eq. S5, and obtain:

$$P(g_1|\,\mathbf{x}) = \frac{1}{\Psi_0\,(\mathbf{x})}\left|\frac{g_1^{n-2}}{x^{n-1}}\right|\;P_\nu\left(\frac{x_1}{g_1}\right)\;\;P_g\left(g_1,\;\frac{g_1}{x_1}x_2,\;\frac{g_1}{x_1}x_3,\;...\;\frac{g_1}{x_1}x_n\right) \tag{S12}$$

For the moments, it is more convenient to start from Eq. S11.

$$\mathrm{E}[g_1|\,\mathbf{x}] = \int_{-\infty}^{+\infty}\mathrm{d}g_1\;g_1\;P(g_1|\,\mathbf{x}) = \frac{1}{P(\mathbf{x})}\int_{-\infty}^{+\infty}\mathrm{d}g_1\,\mathrm{d}\nu\,\frac{g_1}{\nu^{n-1}}\;P_g(g_1,\;\frac{x_2}{\nu},\;...\;\frac{x_n}{\nu})\;P_\nu(\nu)\;\delta(x_1{-}\nu g_1) \tag{S13}$$

We integrate on $g_1$ first, once again operating on the Delta function.

$$\mathrm{E}[g_1|\,\mathbf{x}] = \frac{x_1}{\Psi_0(\mathbf{x})}\int \mathrm{d}\nu\;\frac{1}{\nu^{n+1}}\;P_g\left(\frac{\mathbf{x}}{\nu}\right)P_\nu(\nu) = \frac{x_1}{\Psi_0(\mathbf{x})}\int \mathrm{d}\nu\;P_\nu(\nu)\;\frac{1}{\nu^{n+1}}\;\mathcal{N}\left(\frac{\mathbf{x}}{\nu};\;\mathbf{0},\;\Sigma_g\right) \tag{S14}$$

Using again the definition in Eq. S5, we reach the simple expression:

$$\mathrm{E}[g_1|\,\mathbf{x}] = x_1\frac{\Psi_{-1}(\mathbf{x})}{\Psi_0(\mathbf{x})} \tag{S15}$$

For the second moment, the variance and the Fano factor, the procedure is equivalent.

$$\begin{gathered}\mathrm{E}[g_1^2|\,\mathbf{x}] = x_1^2\frac{\Psi_{-2}(\mathbf{x})}{\Psi_0(\mathbf{x})}\quad;\quad \mathrm{Var}\,[g_1\mid\mathbf{x}] = x_1^2\left[\frac{\Psi_{-2}(\mathbf{x})}{\Psi_0(\mathbf{x})} - \left(\frac{\Psi_{-1}(\mathbf{x})}{\Psi_0(\mathbf{x})}\right)^2\right]\quad;\\ \mathrm{FF}[g_1\,|\,\mathbf{x}] = x_1\left(\frac{\Psi_{-2}(\mathbf{x})}{\Psi_{-1}(\mathbf{x})} - \frac{\Psi_{-1}(\mathbf{x})}{\Psi_0(\mathbf{x})}\right)\end{gathered} \tag{S16}$$

In Section 4 we derive an approximation of these results, as shown in the main test.

#### 3 Statistics of the latent feature for the special case of Rayleigh mixer prior

In the special case $P_\nu(\nu) = \text{Rayleigh}(\alpha)$, Eq. S5 becomes:

$$\Psi_k(\lambda) = \int_0^\infty d\nu\, \nu^{k-n} \frac{\nu}{\alpha^2}\, e^{-\frac{\nu^2}{2\alpha^2}} \frac{e^{-\frac{\lambda}{2\nu^2}}}{\sqrt{(\pi)^n \ \text{Det}\left(C_g\right)}} \quad \text{for} \quad \lambda := \sqrt{\mathbf{x}^\top C_g^{-1}\, \mathbf{x}} \tag{S17}$$

This integral can be solved analytically, using the following general result (Abramowitz and Stegun 2013):

$$\int_0^\infty dy\; y^a\; e^{-b/y^2}\; e^{-c\,y^2} = \left(\frac{b}{c}\right)^{\frac{1+a}{4}} \text{BesselK}_{\frac{1+a}{2}}\left(2\sqrt{b\,c}\right) \tag{S18}$$

After the required substitutions, the result is:

$$\Psi_k(\mathbf{x}) = \frac{1}{\alpha^2} \frac{1}{\sqrt{(2\pi)^n \ \text{Det}\left(\Sigma_g\right)}} (\alpha\,\lambda)^{1+\frac{k-n}{2}}\ \text{BesselK}_{1+\frac{k-n}{2}}\left(\frac{\lambda}{\alpha}\right) \tag{S19}$$

The ratio of two $\Psi_k(\mathbf{x})$ terms takes instead the form:

$$\frac{\Psi_{k_1}(\mathbf{x})}{\Psi_{k_2}(\mathbf{x})} = (\alpha\,\lambda)^{\frac{k_1-k_2}{2}} \frac{\text{BesselK}_{1+\frac{k_1-n}{2}}\left(\frac{\lambda}{\alpha}\right)}{\text{BesselK}_{1+\frac{k_2-n}{2}}\left(\frac{\lambda}{\alpha}\right)} \tag{S20}$$

#### 4 Approximate posteriors for mixer and latent variable

To gain insight about the scaling of the mean and variance, we now consider approximations for the regime of strong input signal:

$$\text{BesselK}_h(z) = \sqrt{\frac{\pi}{2z}} e^{-z} \left[1 + \frac{4h^2-1}{8z} + \frac{(4h^2-1)^2(4h^2-9)}{2(8z)^2} + \mathcal{O}(z^{-3})\right] \quad \text{for} \quad z \gg 1 \tag{S21}$$

The ratio of special Bessel functions then takes the form:

$$\frac{\text{BesselK}_{h_1}(z)}{\text{BesselK}_{h_2}(z)} = 1 + \frac{h_1^2 - h_2^2}{2z} + \mathcal{O}\left(z^{-2}\right) \tag{S22}$$

Replacing the result above into Eq. S20, we obtain:

$$\frac{\Psi_{k_1}(\mathbf{x})}{\Psi_{k_2}(\mathbf{x})} \approx (\alpha\,\lambda)^{\frac{k_1-k_2}{2}} \left[1 + \frac{\alpha}{2\lambda} \left(\left(1 + \frac{k_1 - n}{2}\right)^2 - \left(1 + \frac{k_2 - n}{2}\right)^2\right)\right] \tag{S23}$$

This approximate term be substituted in Eqs. S7, S15 and S16, leading to:

$$\mathrm{E}[\nu|\,\mathbf{x}] = \sqrt{\lambda}\,(1+\mathcal{O}\,(\lambda^{-1})) \tag{S24}$$

$$\mathrm{E}[g_1|\,\mathbf{x}] = \frac{x_1}{\sqrt{\lambda}}\,(1+\mathcal{O}\,(\lambda^{-1})) \quad ; \quad \mathrm{Var}[g_1\,|\,\mathbf{x}] = \left(\frac{x_1}{2\lambda}\right)^2\,(1+\mathcal{O}\,(\lambda^{-1})) \quad ; \\ \mathrm{FF}[g_1\,|\,\mathbf{x}] = \frac{x_1\sqrt{\alpha}}{4\lambda\sqrt{\lambda}}\,(1+\mathcal{O}\,(\lambda^{-1})) \tag{S25}$$

This formulation offers useful insights on how the output of the model depends on both the main drive $x_1$, which is the output of the linear filter applied to the image, and on the global signal, mediated by the $\lambda$ defined in Eq. S17, which roughly corresponds to a norm of **x**. Contextual stimuli leave $x_1$ unvaried, but change $\lambda$, thus scaling down the mean, but also the FF.

Finally, note that in the low-noise case we are considering, when the signal is also small the mean and variance of $g_1$ also asymptote to zero. In our model neuron, this would correspond to zero spontaneous activity and variability, in contrast with cortical data. For this reason in the main text we considered also the model with non-zero noise: it gives similar results to the noiseless model when the signal is large, but better matches cortical data when the signal is small (see Fig. S10).

### 5 Conversion from latent variables to spike counts

To compare the model with neuronal spike counts, we converted the posterior samples of the latent variables into a positive quantity, according to:

$$r = c\sqrt{g_{1+}^2 + g_{1-}^2} \tag{S26}$$

where $1+$ and $1-$ indicate two center, vertically-oriented features, with opposite spatial phases. The choice of phase invariance is purely practical (e.g. reflecting the fact that some awake experiments employ drifting gratings), moreover the form of Eq. S26 preserves all the key findings. In the absence of noise, mean and variance of $r$ can be computed analytically:

$$\mathrm{E}[r|\,\mathbf{x}] = \mathrm{E}\big[c\sqrt{g_{1+}+g_{1-}}|\,\mathbf{x}\big] = c\int_{-\infty}^{+\infty} \mathrm{d}\mathbf{g}\,\sqrt{g_{1+}+g_{1-}}\,P(\mathbf{g}|\mathbf{x}) \tag{S27}$$

This equation can be solved equivalently to Eqs. S13 and S14, with the only difference than integrating over the Delta function will result in the term $\sqrt{x_{1+}^2+x_{1-}^2}$ instead of $x_1$. The same holds for the variance.

$$\mathrm{E}[r|\,\mathbf{x}] = c\sqrt{x_{1+}^2 + x_{1-}^2}\,\frac{\Psi_{-1}(\mathbf{x})}{\Psi_0(\mathbf{x})} \quad \text{and} \quad \mathrm{Var}[r\,|\,\mathbf{x}] = c^2\left(x_{1+}^2 + x_{1-}^2\right)\left[\frac{\Psi_{-2}(\mathbf{x})}{\Psi_0(\mathbf{x})} - \left(\frac{\Psi_{-1}(\mathbf{x})}{\Psi_0(\mathbf{x})}\right)^2\right] \tag{S28}$$
